# Optimal CD8^+^ T-cell memory formation following subcutaneous cytomegalovirus infection requires virus replication but not early dendritic cell responses

**DOI:** 10.1101/2020.12.02.408104

**Authors:** Sandra Dimonte, Silvia Gimeno-Brias, Morgan Marsden, Lucy Chapman, Pragati Sabberwal, Mathew Clement, Ian R Humphreys

**Author notes:** these authors contributed equally (Sandra Dimonte, Silvia Gimeno-Brias). corresponding author, Henry Wellcome Building, Cardiff University, Heath Park, Cardiff, CF14 4XN.

## Abstract

Cytomegalovirus (CMV) induction of large frequencies of highly functional memory T-cells has attracted much interest in the utility of CMV-based vaccine vectors, with exciting preclinical data obtained in models of infectious diseases and cancer. However, pathogenesis of human CMV (HCMV) remains a concern. Attenuated CMV-based vectors, such as replication- or spread-deficient viruses potentially offer an alternative to fully replicating vectors. However, it is not well-understood how CMV attenuation impacts vector immunogenicity, in particularly when administered via relevant routes of immunization such as the skin. Herein we used the murine cytomegalovirus (MCMV) model to investigate the impact of vector attenuation on T-cell memory formation following subcutaneous administration. We found that the spread deficient virus (ΔgL-MCMV) was impaired in its ability to induce memory CD8^+^ T-cells reactive to some (M38, IE1) but not all (IE3) viral antigens. Impaired memory T-cell development was associated with a preferential and pronounced loss of polyfunctional (IFN-γ^+^ TNF-α^+^) T-cells, and was not rescued by increasing the dose of replication-defective MCMV. Finally, whilst vector attenuation reduced dendritic cell (DC) recruitment to skindraining lymph nodes, systematic depletion of multiple DC subsets during acute subcutaneous MCMV infection had a negligible impact on T-cell memory formation, implying that attenuated responses induced by replication-deficient vectors were likely not a consequence of impaired initial DC activation. Thus, overall, these data imply that the choice of antigen and/or cloning strategy of exogenous antigen in combination with the route of immunization may influence the ability of attenuated CMV vectors to induce robust functional T-cell memory.

## Introduction

The beta-herpesvirus human cytomegalovirus (HCMV) is ubiquitous in the human population. Although primary infection is generally asymptomatic in immune-competent individuals, HCMV is a significant cause of morbidity and mortality following congenital infection and primary infection or reactivation in immunocompromised individuals such as immune-suppressed transplant recipients. Despite its pathogenic potential, HCMV has attracted considerable interest as a potential vaccine vector (1,2). HCMV induces unusually high frequencies of antigenspecific CD4^+^ and CD8^+^ T-cells (3) that maintain functionality and gradually increase in abundance over time during viral chronicity (4,5), a process termed ‘memory inflation’ (6). Studies using the rhesus CMV (RhCMV) model have demonstrated that this T-cell response can be redirected towards exogenous antigen with exciting therapeutic potential. Using a CMV-based vector expressing multiple antigens from simian immunodeficiency virus (SIV), induction of broad SIV-specific T-cell responses exhibiting MHC class II and HLA-E restriction was observed (7,8). Importantly, these unusual T-cell responses were accompanied by SIV clearance in approximately half of immunized macaques (7,9), resulting in the consideration of the utility of HCMV-based vaccines in a range of diseases including HIV, tuberculosis (10), Ebola (11) and cancer (12,13).

Despite the potential of CMV-based vaccines, the severity of medical problems associated with HCMV in vulnerable populations remains a major roadblock. Thus, the development of ‘safe’ non-replicating CMV vectors was first explored using the murine CMV (MCMV) model. Whereas temperature-sensitive MCMV failed to induce robust virus-specific CD8^+^ T-cell memory (14), OVA-expressing virus lacking the gene *M94* (MCMV-ΔM94) which is essential for secondary envelopment and cell to cell spread of MCMV (15), induced OVA-specific CD4^+^ and CD8^+^ T-cell responses following systemic challenge (16). Furthermore, Δgl-MCMV lacking the essential viral glycoprotein L (gL) required for viral cell spread induces MCMV-specific CD8^+^ T-cell memory inflation when administered systemically (intra-peritoneally), albeit generally to a lesser degree than replicating virus (17).

Although these studies have indicated excellent safety profiles and induction of effector memory T-cell responses by these attenuated vectors, these investigations have relied upon systemic routes of infection, particularly intra-peritoneal and intra-venous injections. Microenvironmental and cellular differences among anatomic infection sites influence virus-specific T-cell responses (18). Furthermore, the site of infection influences the cell types which viruses infect, transport antigen to draining lymph nodes and present antigen to T-cells (19). Indeed, dendritic cell subsets influence T-cell priming in viral infections by secreting different cytokines leading to Th1 or Th2 dominant responses (20,21) and by preferentially priming CD4^+^ or CD8^+^ T-cells (22).

Injections into the skin are most applicable to vaccination in the clinical setting. Encouraging, preclinical data from the Rhesus macaque model using a replication-competent CMV-based vector was derived from sub-cutaneous immunization (10). However, currently, there is a lack of detailed understanding of how subcutaneous immunization influences the immunogenicity of CMV-based vectors, particularly in the context of attenuated vectors more suitable for broad vaccination strategies. Herein, we used *in vivo* mouse models of infection to perform a systematic investigation of how sub-cutaneous immunization with spread-deficient (Δgl-MCMV) impacts the expansion and function of virus-specific T-cells and the role that relevant DC subsets play to priming of CMV-specific T-cells in this context.

## Materials and Methods

### Mice

BALB/c mice and C57BL/6 mice were purchased from Charles River Laboratories. The following transgenic mouse strains were bred in-house and used for diptheria-toxin (DT)-induced DC depletions. For plasmacytoid dendritic cell (pDC) depletion, BDCA2-DTR (014176-C57BL/6-Tg(CLEC4C-HBEGF)956Cln/J) mice (Jax.org) were maintained as heterozygotes. pDC depletion lasts for 2 to 3 days (23). For Langerhans cell (LC) depletion, Lang-DTREGFP (B6.129S2-Cd207tm3(DTR/GFP)Mal/J) mice (Jax.org) were maintained as homozygotes. Depletion was apparent 24h after DT-injection and lasted for 6–7 days. Complete restoration of the LC pool has been reported after 6 weeks (24). For depletion of cross-presenting dendritic cells, XCR1-DTR mice (Riken B6.Cg-Xcr1<tm2(HBEGF/Venus)Ksho> (RBRC09485)) from Riken (Japan) were maintained as heterozygotes. CD8α^+^ DC depletion is apparent for 4 days and pools are completely restored at day 8 (25). For depletion of classical DCs (cDCs), zDC-DTR (B6(Cg)-Zbtb46tm1(HBEGF)Mnz/J) mice (Jax.org) were maintained as homozygotes. In bone marrow chimeras, CD11c^+^-DCs were depleted at 12h post DT injection. Reconstitution starts at day 5 and pools are completely restored in the spleen at day 7 (26). To generate chimeric mice, C57BL/6 mice were irradiated (2 x 550rad) and transfused with 2×10^6^ zDC-DTR or, in some experiments, XCR1-DTR bone marrow cells 24h post-irradiation. Mice were subsequently treated with antibiotic-supplement water (Baytril, Bayer) for 4 weeks. DC depletion and MCMV infections were performed 6-8 weeks post-irradiation. DT was injected intraperitoneally (i.p.) at 300ng/mouse on day −1 prior to viral infection or, in XCR1-DTR mice, at day −3/−2 and day-1. Where non-chimeric, heterozygote mice were used (BDCA2-DTR), both transgenic and control mice without transgene were injected with DT. In the case of homozygote or chimeric mice (Lang-DTREGFP, XCR1-DTR, zDC-DTR), control groups were injected with PBS.

### Viral infections

Mice were injected subcutaneously (s.c.) in the sternum area with 2×10^5^ PFU WT-MCMV (BAC-derived MCMV strain K181 (a kind gift from Alec Redwood, University of Western Australia) in 50μl sterile PBS or with replication deficient ΔgL-MCMV/K181(17) (a kind gift from Chris Snyder, Thomas Jefferson University) at 2×10^5^ PFU, 8×10^5^ PFU or 2×10^6^ PFU in 50μl sterile PBS. ΔgL-MCMV was grown and titred using 3T3 cells that express glycoprotein L (gL). Mice were sacrificed 24h or 16-18 or 43 weeks post-infection, as indicated in figure legends.

### Leukocyte isolation and analysis

Leukocytes were isolated from spleen or axillary draining lymph nodes as described previously (27). Peripheral blood was collected from the lateral tail vein into heparin-coated tubes (Sarstedt) at time points stated in the legends. Depletion of Langerhans cells (LCs) was confirmed by a migration assay from ear skin using mouse 6Ckine, as described elsewhere (28). In brief, ears were removed and dorsal skin sheet was isolated via peeling from the ear cartilage using tweezers. The dorsal skin sheets where placed ventral side down in RPMI 1640 (Gibco) containing 10% FCS for 2-4h at 37°C to release non-DCs. The skin sheets were then transferred into another 1ml medium containing 0.1 μg of recombinant mouse 6Ckine (BioLegend) to enhance DC-migration. After 24h, skin sheets were again transferred into 1ml fresh medium containing 6Ckine and incubated for a further 24h. Supernatants from both time-points were pooled and assessed for migrating DCs by flow cytometry.

For DC analysis, cells were stained with LIVE/DEAD Zombie Aqua dye (BioLegend) and incubated with FC block and subsequently stained with a combination of the following anti-mouse antibodies (all from BioLegend, eBioscience, or BD Biosciences): anti-CD8α (53-6.7), anti-CD11c (N418), anti-CD11b (M1/70), anti-IAIE (M5/114.15.2), anti-Siglec H (551), anti-CD103 (2E7), anti-CD80 (16-10A1), antiCD86 (GL-1), anti-CD205 (NLDC-145), anti-EpCAM1 (G8.8), and anti-4-1BBL (TKS-1).

For detection and phenotypical analysis of MCMV-specific CD8^+^ T-cells, leukocytes were stained with LIVE/DEAD Zombie Aqua dye (BioLegend) and incubated with FC block (BioLegend), followed by tetramer incubation (25μg/ml) for 15min at 37°C. Biotinylated monomers loaded with H-2Kb restricted M38 (SSPPMFRV) and IE3 (RALEYKNL) peptides were generated by the NIH tetramer facility (Emory, USA), and were conjugated using PE-labelled streptavidin (BioLegend) at a 4:1 molar ratio to generate tetramers (29). Subsequently, a combination of the following anti-mouse antibodies was used to stain for surface markers (all from BioLegend, eBioscience, or BD Biosciences): anti-CD3ε (145-2C11), anti-CD4 (RM4-5), anti-CD8α (53-6.7), anti-CD44 (IM7), anti-CD62L (MEL-14), anti-CD11a (M17/4), anti-CD69 (H1.2F3), anti-CD103 (2E7), anti-CD127 (A7R34), anti-KLRG1 (MAFA).

For functional analysis, leukocytes were stimulated for 6h at 37°C with peptide M38 (M38, IE3) at a final concentration of 2μg/ml in the presence of brefeldin A (1μg/ml) (BD Biosciences). Cells were then stained with LIVE/DEAD Zombie Aqua dye (BioLegend) and incubated with FC block. Cells were subsequently stained with anti-CD8 (53-6.7), and cells were then fixed in 4% paraformaldehyde and permeabilized with saponin buffer (PBS, 2% FCS, 0.05% sodium azide, and 0.5% saponin) in order to perform intracellular cytokine staining using anti-TNαz (MP6-XT22) and anti-IFNγ (XMG1.2). All data were acquired using Attune NxT Flow Cytometer from Thermo Fisher.

### Statistics

Paired data were evaluated for statistical significance using non-parametric Mann-Whitney U test in Prism 8.2.1 for macOS (GraphPad). Where multiple comparisons were performed, a 1-way ANOVA test was used for single time-point multi-group analysis and a 2-way ANOVA was used for time course data.

### Ethics

All mouse experiments were conducted according to U.K. Home Office guidelines under a Home Office project license (PPL P7867DADD, granted to I. R. Humphreys) at the Home Office-designated facility at Heath Park, Cardiff University.

## Results

### MCMV replication is required for optimal T-cell memory induction following subcutaneous infection

We first sought to investigate the immunogenicity of ΔgL-MCMV following subcutaneous infection. C57BL/6 mice were administered 2×10^5^ PFU ΔgL-MCMV or ‘WT’-MCMV (pARK25), and peripheral blood CD8^+^ T-cell responses to the ‘inflationary’ H2K^b^-restricted MHC class I-restricted peptides M38 and IE3 were measured over time. ΔgL-MCMV induced significantly lower frequencies of IFN-γ secreting IE3 and M38-specific CD8^+^ T-cells in peripheral blood compared to WT-MCMV during the acute phase of the response 7 days p.i (Fig. 1A & B). Interestingly, during the memory phase (from 4 weeks p.i), only M38-specific IFN-γ producing T-cells remained significantly lower in mice infected with ΔgL-MCMV as compared to WT-MCMV (Fig. 1A & B). Furthermore, when tissue responses were measured in the spleen at 19 weeks p.i, both peptide stimulation (Fig. 1C & D) and tetramer staining (Fig. 1E, F & G) of splenocytes demonstrated significantly reduced frequencies and numbers of M38-specific but not IE3-specific T-cells in ΔgL-MCMV infected mice as compared to WT-MCMV.

**Figure 1:**
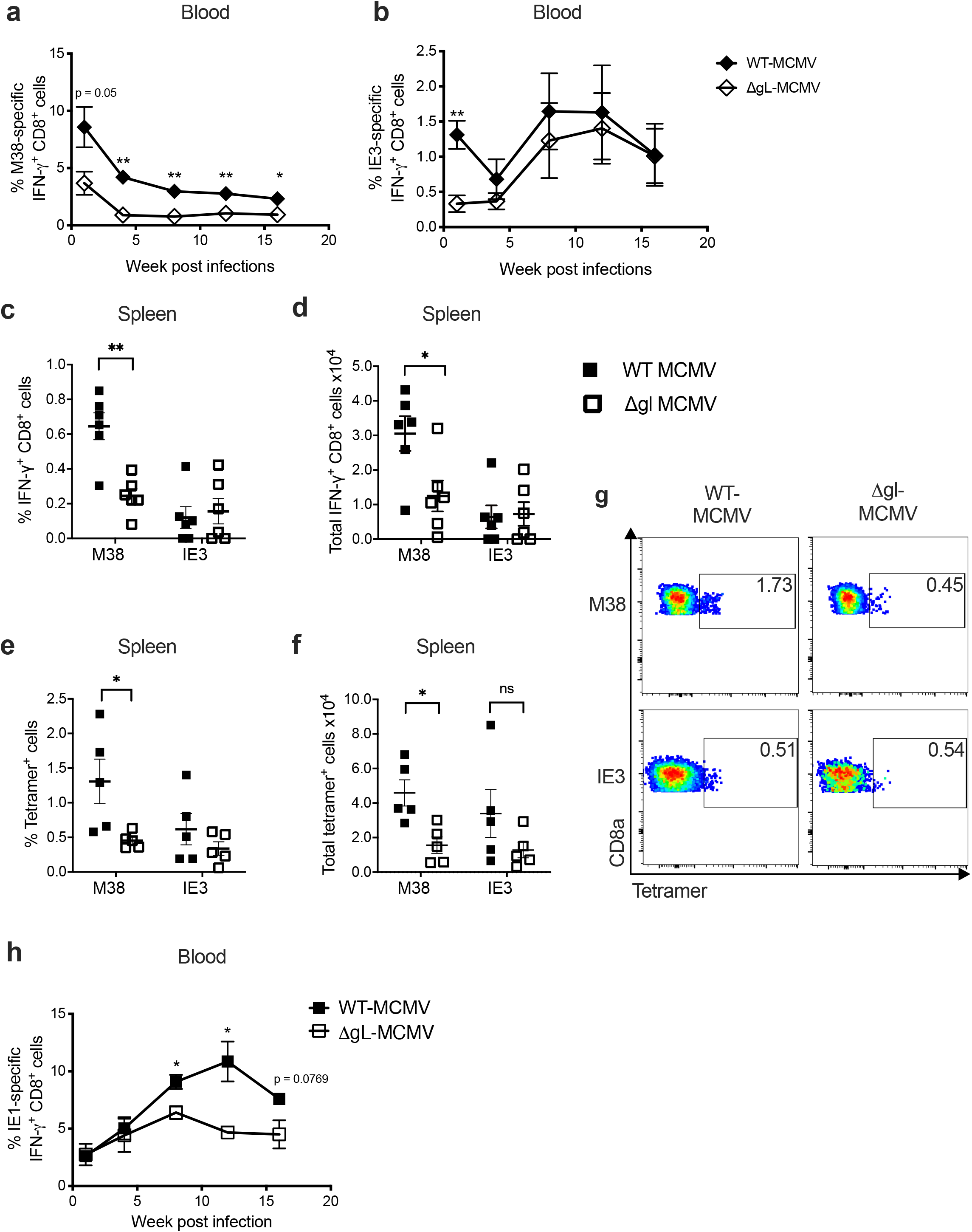
Subcutaneous infection with ΔgL□MCMV results in impaired memory T-cell induction in C57BL/6 and BALB/c mice. **A-B**: C57BL/6 mice were administered s.c. with 2×10^5^ PFU of WT-MCMV or Δgl. MCMV Leukocytes were isolated from peripheral blood on week 1, 4, 8, 12 and 16. Frequencies and total numbers or functional (CD8^+^IFNγ^+^) T-cells specific for IE3 **(A)** or M38 **(B)** were quantified using flow cytometry. Data shown as mean ± SEM (n = 4) **C-G**: Leukocytes were isolated from spleens at week 19 post infection (p.i) and quantified using flow cytometry. Data shown as mean ± SEM (n = 6 mice per group). **C**: frequencies of functional (CD8^+^IFN-γ’) T-cells specific for M38 or IE3 **D:** Total numbers of functional (CD8^+^IFN-γ^+^) T-cells specific for M38 or IE3. **E:** Frequencies of tetramer-binding CD8^+^ T-cells specific for M38 or IE3. **F:** Total numbers of tetramer-binding CD8^+^ T-cells specific for M38 or IE3. **G:** Representative flow cytometry plots of frequencies of tetramer-binding CD8^+^ T-cells specific for M38 or IE3 are shown. **H:** BALB/c mice were infected/immunised s.c. with 2×10^5^ PFU WT-MCMV or ΔgL□MCMV Leukocytes were isolated from peripheral blood on week 1, 4, 8, 12 and 16 p.i. Frequencies of pp89-specific functional T-cells (CD8^+^IFNγ^+^) were quantified using flow cytometry. Data shown as mean ± SEM (n = 3-4), representing for 2 experiments. *p ≤ 0.05, **p ≤ 0.01.

We further investigated the epitope-specific impact of MCMV attenuation on virusspecific memory T-cell formation by studying another inflationary CD8^+^ T-cell population, namely IE1-specific T-cells in BALB/c mice (6). BALB/c mice immunized with ΔgL-MCMV or WT-MCMV induced similar frequencies of IE1-specific IFN-γ secreting CD8^+^ T-cells during the acute phase of infection. However, at week 4 p.i, these responses diverged, with reduced memory inflation in ΔgL-MCMV infected mice throughout the time-course of infection (Fig. 1H). Thus, overall, these data suggest that ΔgL-MCMV is impaired in its ability to induce memory T-cell formation following subcutaneous infection, although the impact may vary depending on the antigen in question.

### T-cell memory induced by ΔgL-MCMV is functionally and phenotypically distinct from T-cells induced by replication-competent MCMV

Next, we assessed whether MCMV replication impacted cytokine expression of inflationary CD8^+^ T-cell responses, focusing on the robust replication-dependent M38-specific responses. After 18 weeks p.i., most M38-specific T-cells in peripheral blood and spleens of C57BL/6 mice were polyfunctional (IFN-γ^+^TNF-α^+^) in WT-MCMV infection (Fig. 2A & B). Strikingly, we observed a substantial reduction in polyfunctional cells in peripheral blood in mice subcutaneously injected with ΔgL-MCMV. Although we observed a slight (not significant) increase in single positive cells (IFN-γ^+^TNF-α^-^ or IFN-γ^-^TNF-α^+^) in blood, analysis of these responses in spleens revealed a preferential loss of polyfunctional cells with no evidence of conversion to single IFN-γ or TNF-α producers. In addition, proportional compositions of the response were analyzed and revealed that, especially in the blood, a lower proportion of M38-specific polyfunctional cells was found in mice infected with ΔgL-MCMV (Fig. 2D & E). Thus, ΔgL-MCMV infection via the sub-cutaneous route leads to a preferential impairment of MCMV-specific polyfunctional memory T-cell formation.

**Figure 2:**
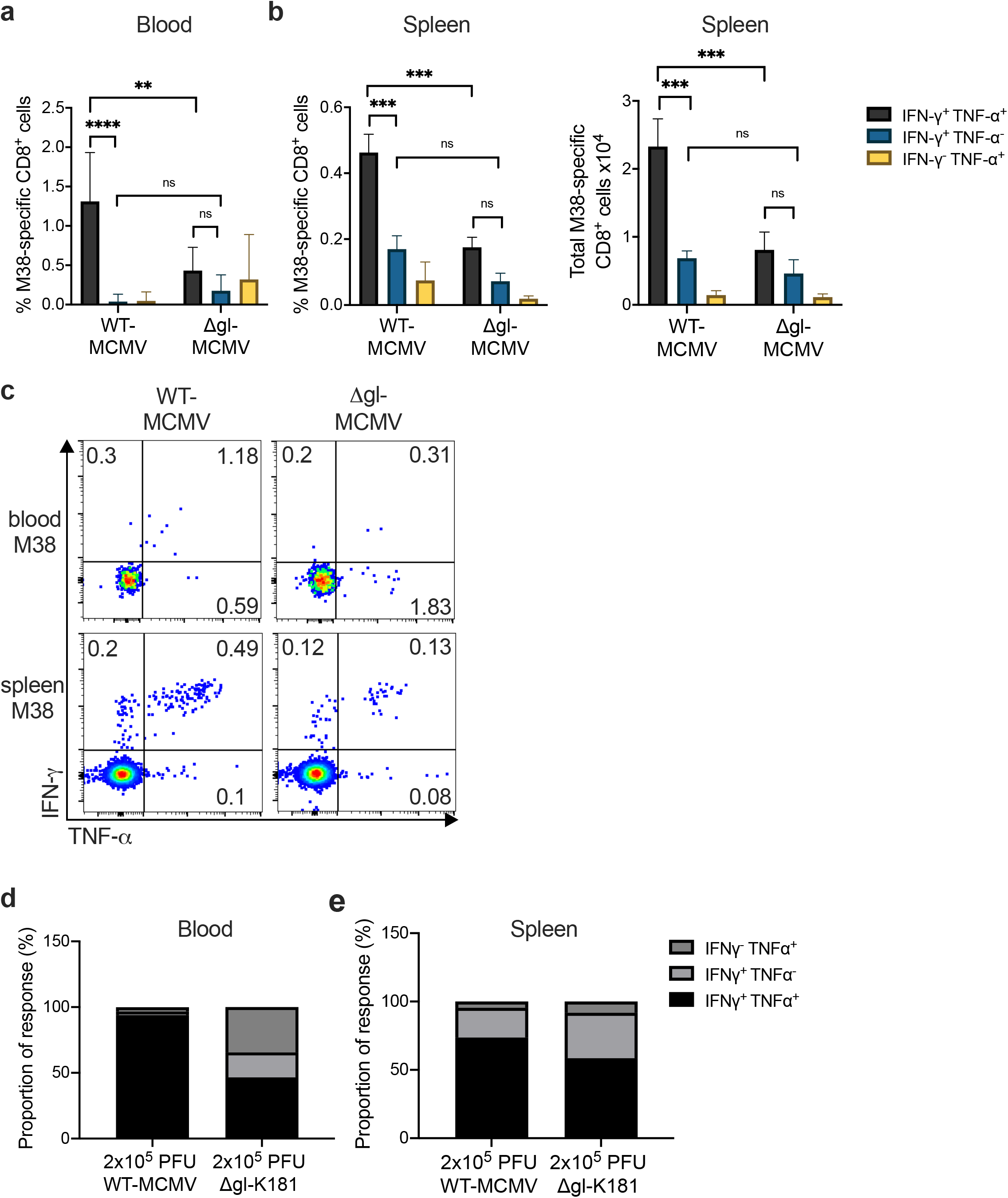
Subcutaneous infection with ΔgL□MCMV leads to a preferential loss of polyfunctional memory CD8^+^ T-cells. C57BL/6 mice were infected s.c. with 2×10^5^ PFU of WT-MCMV or ΔgL-MCMV. Leukocytes were isolated from peripheral blood on week 18 p.i. and spleens on week 19 pi and quantified using flow cytometry. Data shown as mean ± SEM (n=6). **A:** Frequencies of polyfunctional (IFN-γ^+^TNF-α^+^) and monofunctional (IFN-γ^+^TNF-α^-^ or IFN-γ^-^TNF-α^+^) T-cells in peripheral blood, gated on live CD8^+^ cells specific for M38. **B:** Frequencies and total numbers of polyfunctional and monofunctional T-cells in spleens, gated on live CD8^+^ cells specific for M38. **C:** Representative flow cytometry plots of polyfunctional and monofunctional T-cells in spleen and blood of mice infected with WT-MCMV vs. ΔgL□MCMV **D:** proportion of M38-specific functional response. The sum of total numbers of polyfunctional and monofunctional T-cells in spleens equals 100%. Proportions of poly- and monofunctional subsets have been calculated at proportions of 100%. **E**: proportion of IE3-specific functional response. The sum of total numbers of polyfunctional and monofunctional T-cells in spleens = 100%. Proportions of poly- and monofunctional subsets have been calculated as proportions of 100%. \**p* ≥ 0.05, \*\**p* ≤ 0.01, \*\*\**p*≤ 0.001, \*\*\*\**p* ≤ 0.0001

Next, phenotypic analysis of M38-specific CD8^+^ T-cells produced upon s.c. ΔgL-MCMV infection was investigated. Proportions of splenic M38-specific central memory T-cells (TCM) and effector memory T-cells (TEM) (assessed using CD62L and CD44 expression) were comparable after ΔgL-MCMV and WT-MCMV infection with, as expected, most cells exhibiting effector memory phenotype (Fig. 3A). However, in accordance with reduced total numbers of M38-specific CD8^+^ T-cell responses, significant reduction in TCM, and TEM in ΔgL-MCMV-infected mice were observed (Fig. 3B). Furthermore, when expression of KLRG1 and CD127 on CD44^+^-cells was measured, we noted a significant reduction of total numbers of both memory-precursor effector cells (MPECs: CD127^+^KLRG1^-^) and short-lived effector cells (SLECs: CD127^-^ KLRG1^+^) in ΔgL-MCMV infected mice (Fig. 3C-E). Collectively, these data suggest that ΔgL-MCMV induces fewer numbers of M38-specific memory CD8^+^ T-cells without preferentially impacting a particular cell subset(s).

**Figure 3:**
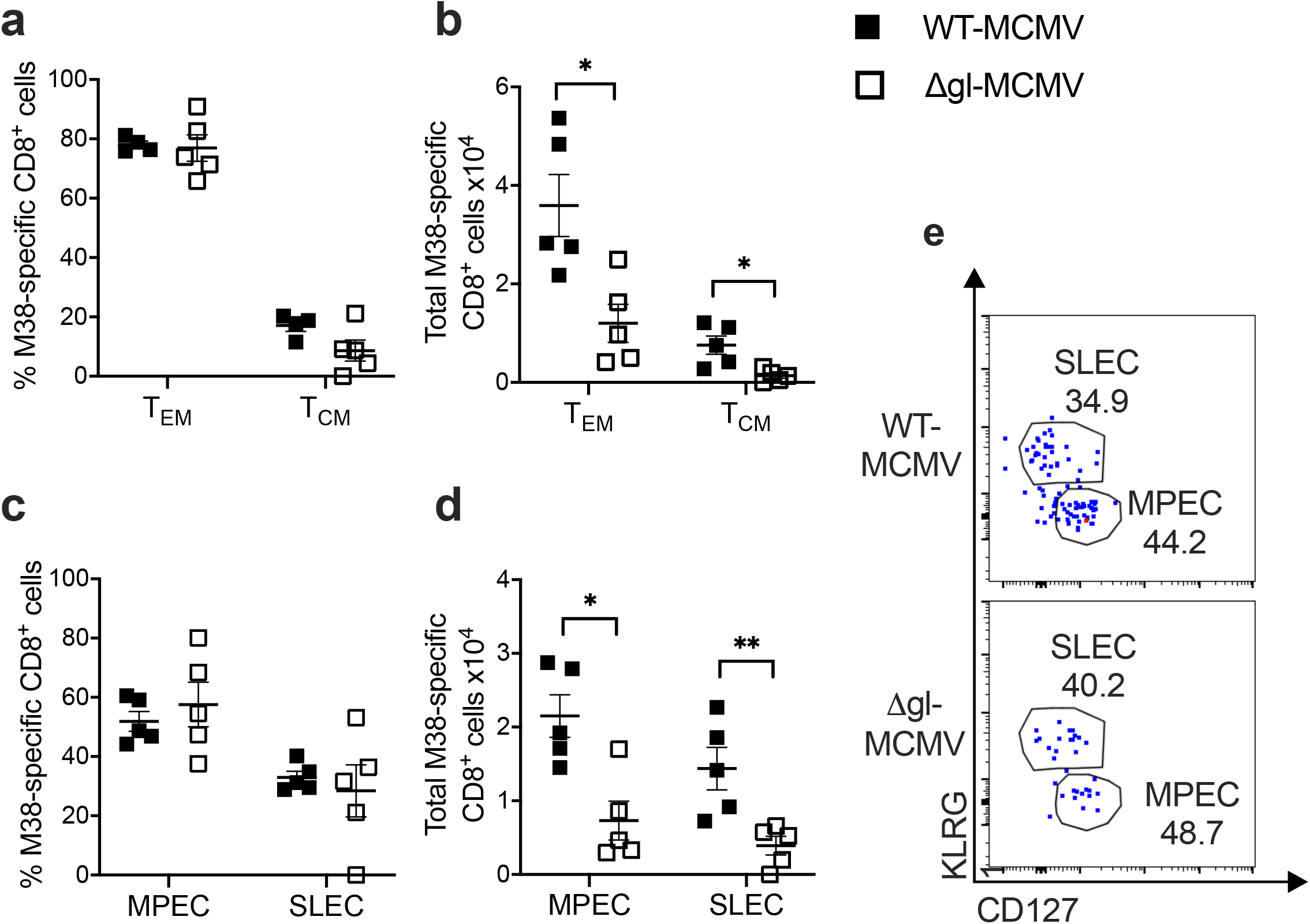
Virus replication is required for maximal formation of multiple memory CD8^+^ T-cell subsets following subcutaneous infection. C57BL/6 mice were infected s.c. with 2×10^5^ PFU of WT-MCMV or ΔgL□MCMV Leukocytes were isolated from spleens at week 19 pi and quantified using flow cytometry. Data shown as mean ± SEM (n=6). **A:** Frequencies of M38 tetramer-binding effector memory T-cells (T_EM_: CD8^+^ CD44^+^ CD62L^-^), central memory T-cells (T_CM_: CD8^+^ CD44^+^ CD62L^+^). **B:** Total numbers of M38 tetramer-binding effector memory T-cells (T_EM_: CD8^+^ CD44^+^ CD62L^-^), central memory T-cells (T_CM_: CD8^+^ CD44^+^ CD62L^+^). **C:** Frequencies of MPECs (CD127^+^KLRG1^+^) and SLECs (CD127^-^KLRG1^+^). **D:** Total numbers of MPECs (CD127^+^KLRG1^+^) and SLECs (CD127^-^KLRG1^+^). **E:** Representative flow cytometry plots of MPECs (CD127^+^KLRG1^+^) and SLECs (CD127^-^KLRG1^+^) specific for M38 in mice infected with WT-MCMV or ΔgL□MCMV. p ≤ 0.05, **p ≤ 0.01, ***p ≤ 0.001

### Increasing dosing of ΔgL-MCMV does not augment T-cell memory formation

Impaired induction of functional T-cell memory by ΔgL-MCMV may simply reflect the requirement for more virus particles in the initial infection and thus, in a vaccination setting, could be overcome by using a larger dose of replication-deficient MCMV. To assess this, mice were infected subcutaneously with 2×10^5^ PFU, 8×10^5^ PFU or 2×10^6^ PFU ΔgL-MCMV or, as control, 2×10^5^ PFU WT-MCMV. Increasing the dose of ΔgL-MCMV as much as ten-fold had only a negligible impact on M38 specific CD8^+^ T-cell memory accumulation 43 weeks later (Fig. 4A). Functional analysis of T-cells showed a trend towards fewer polyfunctional (IFN-γ^+^TNF-α^+^) T-cells in all ΔgL-MCMV doses as compared to WT-MCMV (Fig. 4D & E), with no discernible increase in frequencies of polyfunctional T-cells after increasing the inoculum of ΔgL-MCMV. Overall, these data suggest that impaired memory T-cell formation following infection with replication-deficient MCMV cannot be easily overcome by increasing the initial inoculum.

**Figure 4:**
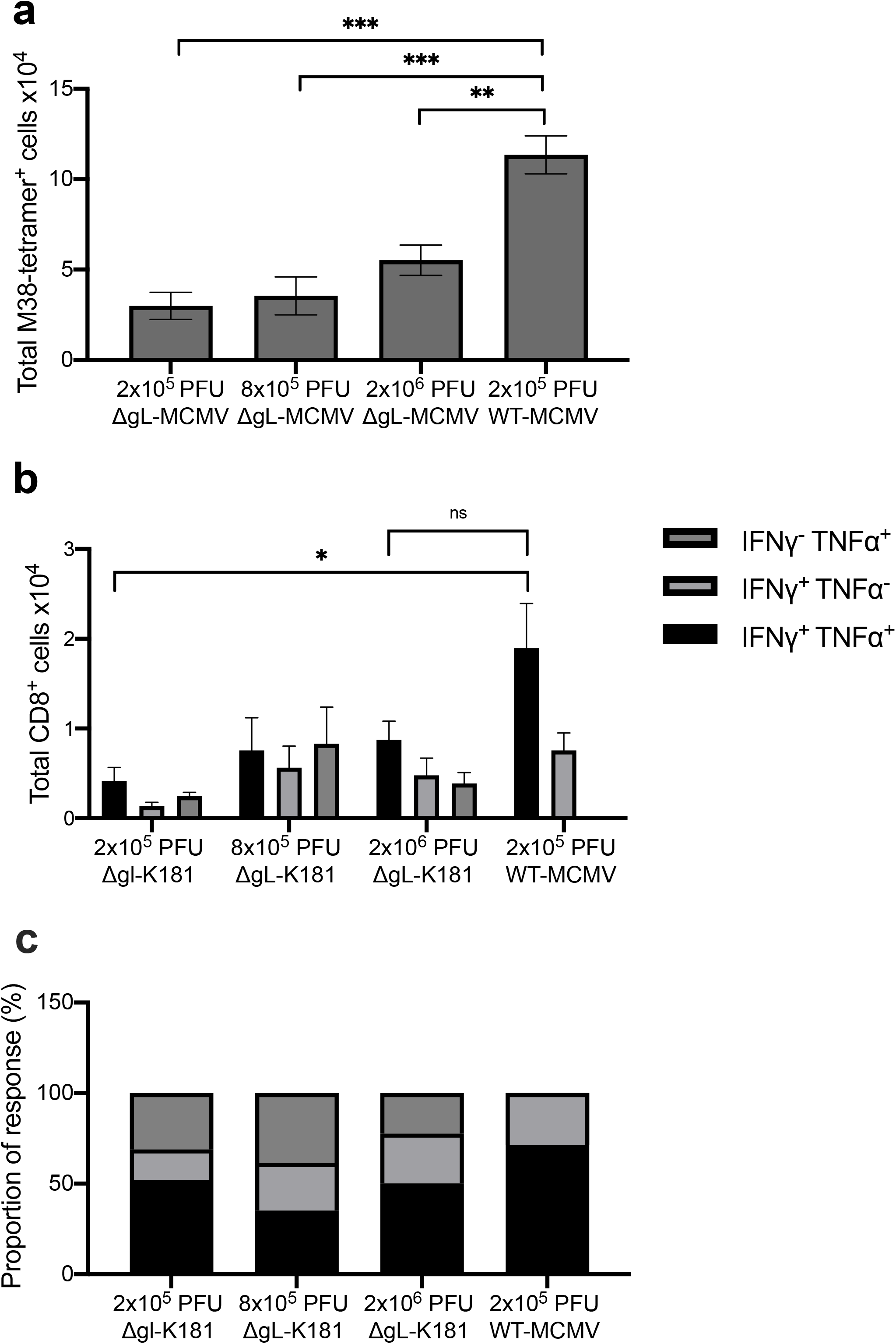
Increasing dosing of ΔgL-MCMV does not augment memory T-cell formation. C57BL/6 mice were infected s.c. with 2×10^5^ PFU (n=4), 8×10^5^ PFU (n=5) or 2×10^6^ PFU (n=5) of ΔgL-MCMV or 2×10^5^ PFU WT-MCMV (n=6). Leukocytes were isolated from spleens 43 weeks p.i. and quantified using flow cytometry. **A:** Total numbers of tetramer binding CD8^+^ T-cells specific for M38. **B:** total numbers of polyfunctional (IFN-γ^+^TNF-α^+^) and monofunctional (IFN-γ^+^TNF-α^-^ or IFN-γ^-^TNF-α^+^) T-cells in spleens, gated on live CD8^+^ cells specific for M38. **C:** Proportion of M38 specific functional response. The sum of total numbers of polyfunctional and monofunctional T-cells in spleens equals 100%. Proportions of poly-and monofunctional subsets have been calculated at proportions of 100%. Data shown as mean + SEM. *p ≤ 0.05, **p ≤ 0.01, ***p ≤ 0.001

### Initial T-cell priming by dendritic cells has minimal impact on inflationary T-cell memory formation after subcutaneous MCMV infection

We have previously demonstrated that manipulation of cytokine signaling during initial MCMV infection and T-cell priming can influence memory T-cell inflation (27,30) and can be manipulated to enhance immunogenicity of replication-deficient MCMV (25). Given that reduced memory T-cell formation following sub-cutaneous ΔgL-MCMV infection was accompanied by impaired responses during the acute phase of infection, we hypothesized that reduced memory T-cell formation in these mice may reflect defective T-cell priming. Thus, we measured cDC responses in skin-draining axillary lymph nodes that we confirmed had drained from the injection site used, using Evans blue injection. Following ΔgL-MCMV infection, we observed reduced classical DC (cDC, CD11c^+^MHCII^hi^) accumulation in draining lymph nodes 24h after ΔgL-MCMV infection, as compared to WT MCMV (Figure 5A), suggesting a reduction in global DC responses may contribute to reduced memory T-cell formation induced by ΔgL-MCMV. To test this and provide improved clarity of the DC subsets that contribute to MCMV-induced memory T-cell formation, we systematically depleted DCs using established transgenic mice that enable diptheria toxin-mediated transient depletion of either pDCs, cross-presenting (XCR1^+^) DCs, classical DCs and Langerhans cells. Published methodology demonstrated that DC depletion lasts for at least 3-7 days (23–26) and is thus removing DCs during the T-cell priming in acute infection, which we confirmed in-house (Supplemental Fig. 1). As depletion of classical DCs in zDC-DTR mice is fatal (26), bone marrow chimeras of WT C57BL/6 mice were generated prior to cDC-depletion. The same approach was used with XCR1-DTR mice for the depletion of cross-presenting DCs to ensure comparability when assessing the two overlapping DC populations.

**Figure 5:**
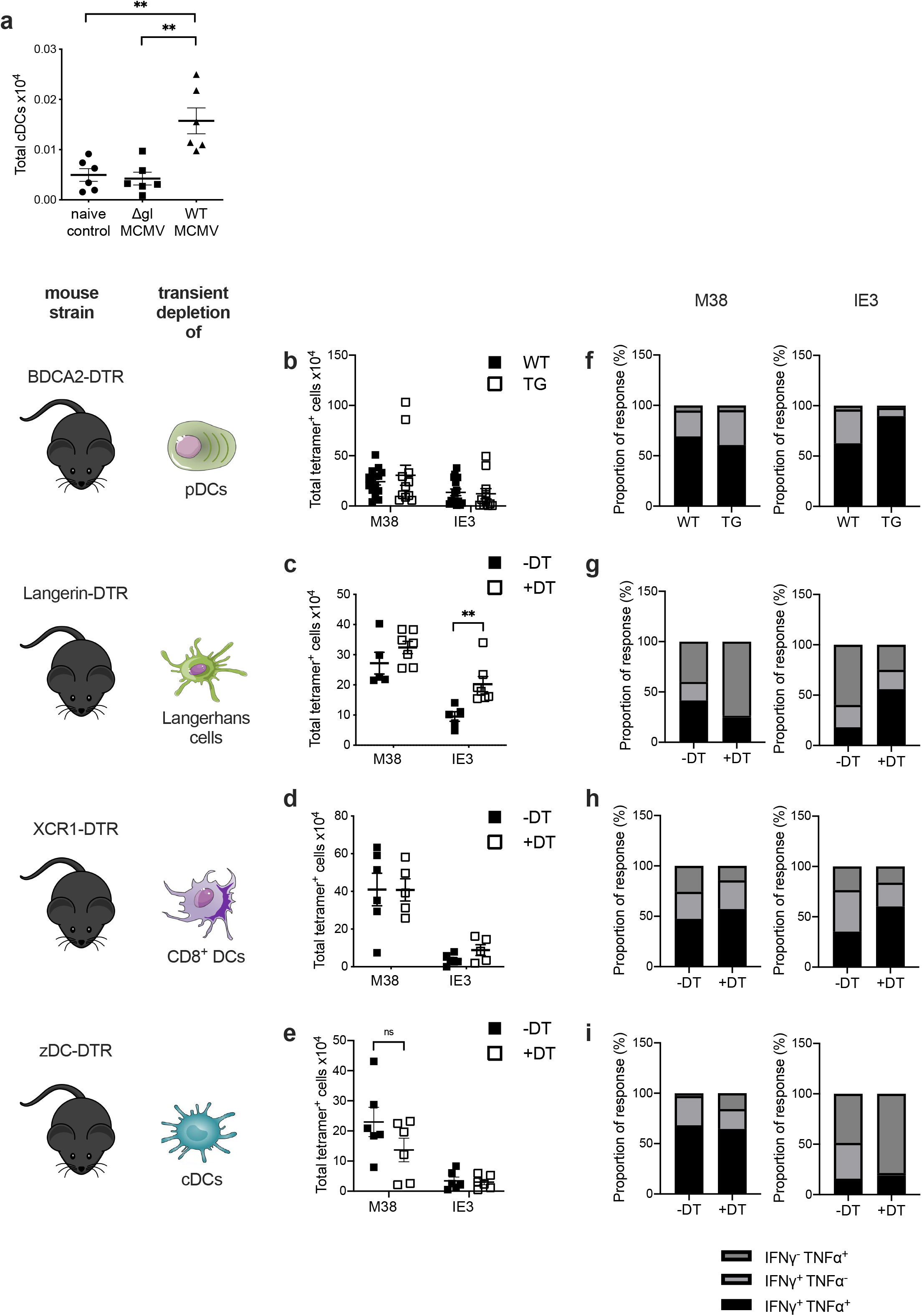
Depleting DC subsets during initial subcutaneous ΔgL-MCMV infection does not dramatically impact memory T-cell formation. **A:** Total numbers of cDCs (CD11c^+^ MHCII^+^) isolated from axillary lymph nodes 24h post s.c. infection were quantified using flow cytometry. **B-E:** total numbers of tetramer binding CD8^+^ T-cells specific for M38 or IE3 in spleens quantified by flow cytometry. **F-I:** proportion of M38-and IE3 specific functional response. The sum of total numbers of M38- or IE3-specific polyfunctional (IFN-γ^+^TNF-α^+^) and monofunctional (IFN-γ^+^TNF-α^-^ or IFN-γ^-^TNF-α^+^) T-cells in spleens = 100%. Proportions of poly-and monofunctional subsets have been calculated as proportions of 100%. **B:** BDCA2-DTR transgenic (TG) and non-transgenic (WT) mice were injected with 300ng diphtheria toxin (DT) 24 hours before s.c. infection with 2×10^5^ PFU WT-MCMV. Leukocytes were isolated from spleens at 19/18 weeks pi. Data shown as mean ± SEM (n= 14 WT and 12 TG) representative for 2 experiments. **C:** XCR1-DTR transgenic (TG) and non-transgenic (WT) mice were injected with 300ng diphtheria toxin (DT) 72 hours and 24 hours before s.c. infection with 2×10^5^ PFU WT-MCMV. Leukocytes were isolated from spleens at 18 weeks pi. Data shown as mean ± SEM (n= 6 WT and 5 TG). **D:** LANG-DTREGFP mice were injected with 300ng diphtheria toxin (+DT) or not (-DT) 24 hours before s.c. infection with 2×10^5^ PFU WT-MCMV. Leukocytes were isolated from spleens at 18 weeks pi. Data shown as mean ± SEM (n= 10 +DT and 7-DT). **E:** zDC-DTR bone marrow chimeras were injected with 300ng diphtheria toxin (+DT) or not (-DT) 24hours before s.c. infection with 2×10^5^ PFU WT-MCMV. Leukocytes were isolated from spleens 19 weeks pi. Data shown as mean ± SEM (n= 5 +DT and 6-DT). *p ≤ 0.05, **p ≤ 0.01

We observed no impairment of memory T-cell formation after subcutaneous MCMV infection after depletion of pDCs (Fig. 5B), Langerhans cells (Fig. 5C) or, in contrast to data derived from mice with genetic defects in cross-presenting DCs (and thus throughout the time-course of infection (31)), XCR1^+^ DCs (Fig. 5D). Following depletion of classical DCs (including cross-presenting DCs) using zDC mice, we noted a trend in impaired M38-but not IE3-specific CD8^+^ T-cell memory development after robust DC depletion (Fig. S1), although differences were not statistically significant (Fig. 5E). We also assessed the influence of early DC responses on memory T-cell functionality. We observed that after depletion of pDCs, Langerhans cells and XCR1^+^ DCs (Fig. 5F-H), memory CD8^+^ T-cell polyfunctionality was unaltered (Fig. 5I). Interestingly, in the case of Langerhans cell depletion, we observed an increased accumulation of IE3-specific T-cells after depletion (Fig. 5C), consistent with known suppressive functions of Langerhans cells (32). Thus, these data argue against a major influence for early DC activation in the immunogenicity of replicating MCMV. Overall, our data imply that strategies beyond increasing dosing and targeting DC activation should be considered when attempting to improve the immunogenicity of subcutaneous vaccination with replication-deficient CMV vectors.

## Discussion

In this study, we demonstrated that subcutaneous infection with spread-deficient MCMV induces significantly inferior T-cell memory responses to inflationary epitopes when administered subcutaneously. Previous detailed investigations of the impact of MCMV replication on virus-induced T-cell responses focused primarily on a systemic infection via the peritoneum where ΔgL-MCMV induced memory T-cell responses, albeit at lower frequencies as compared to replicating virus. The same study noted that footpad infection of ΔgL-MCMV failed to induce notable memory responses (17). We now show that subcutaneous infection with ΔgL-MCMV induces inferior polyfunctional memory T-cell responses as compared to replicating MCMV, which corresponds to a reduction in viral-specific memory T cell subsets.

Why ΔgL-MCMV induces inferior memory T-cell responses following subcutaneous infection is unclear. Studies using replication deficient Δm94-MCMV have reported that latent load of spread-deficient virus is comparable to that of replicating virus (16), suggesting that impaired memory induction is not a consequence of reduced latent virus that leads to CMV reactivation and antigen presentation required for maintenance of long-term T-cell memory (33,34). Furthermore, merely increasing the dose of ΔgL-MCMV did not improve induction of memory formation, suggesting that the initial burst of replication and the virions produced also did not greatly influence memory T-cell formation. Cell-to-cell spread is a key feature of HCMV replication and viral infection of DCs (35). Whether this route of cell spread, lacking in ΔgL-MCMV infection, induces immune activation relevant for induction of memory T-cell formation is currently unclear.

We noted that subcutaneous infection with ΔgL-MCMV induced the accumulation of fewer classical DCs into draining lymph nodes during acute infection, and this correlated with reduced T-cell responses during this phase, at least in the C57BL/6 model. This contrasts with data obtained in the intraperitoneal infection model where non-replicating MCMV actually enhanced initial DC accumulation by triggering less type I IFN that drives DC apoptosis in this model. Consequently, the authors observed elevated T-cell responses during the acute phase of non-replicating MCMV infection (36). Similarly, in BALB/c mice, Dalod and colleagues demonstrated that type I IFN ablated splenic cDCs responses, with a subsequent delay of acute CD8^+^ T-cell responses (37). Differences between these observations and our study may reflect differential cell tropism of the virus, or altered induction by MCMV of type I IFN following systemic versus subcutaneous infection. Irrespective, these differences highlight the importance of the consideration of route of infection when considering early immunological events during CMV-based vector immunization.

Despite the impact of MCMV replication on DC accumulation in lymph nodes, systematic depletion of different DC subsets during the first few days of infection revealed no obvious role for pDC, Langerhans cells or XCR1^+^ DCs in facilitating inflationary memory T-cell formation. The latter observation is in accordance with data derived from *Batf^-/-^* mice that lack cross-presenting DCs, demonstrating that the absence of these cells throughout the infection time-course had a minimal impact on memory inflation despite impacting T-cell priming (31). Transient depletion of all cDCs (including cross-presenting cells) showed a trend in reduced M38-but not IE3-specific CD8^+^ memory T-cells. However, these data argue against a dominant early role for DCs in determining memory T-cell formation. In fitting with this, we failed to boost M38- and IE3-specific memory CD8^+^ T-cell responses induced by subcutaneous ΔgL-MCMV infection co-administered with activating ligands of DC-expressed endosomal TLR7 or TLR9 (data not shown). This contrasts the improved memory T-cell formation when using the alarmin IL-33 (30), that is released upon cell damage after viral infections (38), as an adjuvant. Similarly, blocking the function of the immune-suppressive cytokine IL-10 during initial MCMV infection increases memory T-cell formation, thus demonstrating that early immunological events can be manipulated to increase CMV-induced memory T-cell formation. However, in line with the established role for non-hematopoietic cells in sustaining memory T-cell inflation (39,40). these data imply that DCs may not represent useful targets for strategies that aim to improve immunogenicity of replication-deficient CMV vectors.

We noted an epitope specific impact of virus replication on memory T-cell induction after subcutaneous infection with ΔgL-MCMV, with an impairment of IE1-specific responses in BALB/c mice, and M38 but not IE3-specific responses in C57BL/6 mice. Differential dependency of MCMV-specific CD8^+^ T-cell memory on CD4^+^ T-cell help has been described in C57BL/6 mice (41). However, given that IE3-specific responses are preferentially impacted by CD4^+^ T-cell deficiency, the observation that IE3-specific inflationary responses were unaffected by replication-deficient MCMV argues against this hypothesis. Moreover, both IE3- and M38-specific T-cell responses during the acute phase of infection were impacted by replication deficiency. Understanding why antigen-specific memory T-cell responses are differentially affected by virus replication following infection via the skin may inform the design of CMV vectors in the future.

One way to improve memory induction by a replicating vector could be simply to increase the infectious dose and initial antigen exposure, triggering a greater primary T-cell response and thereby possibly increasing the formation of a memory T-cell pool. Indeed, careful dose titration experiments using replicating MCMV clearly show a positive correlation between virus dose and amplitude of virus-specific memory induction (42). In our study however, we observed that increasing the dose of ΔgL-MCMV had little impact on memory T-cell formation and polyfunctionality, certainly at the doses used. Whether the amount of antigen is already maximal, or whether any beneficial effects of increasing antigen load are counteracted by an elevated type I IFN responses and subsequent DC apoptosis are unclear. Irrespective, collectively our data suggest that alternate approaches beyond dose elevation and DC manipulation may be needed to maximize immunogenicity of CMV-based vaccine vectors for safe and efficacious utility in the future.

## List of abbreviations

CMV: Cytomegalovirus
HCMV: human CMV
MCMV: murine CMV
RhCMV: rhesus CMV
DC: dendritic cell
IFN-γ: Interferon γ
TNF-α: Tumor necrosis factor α
SIV: simian immunodeficiency virus
HIV: human immunodeficiency virus
IE3: Immediate early 3
IE1: Immediate Early 1
CD: cluster of differentiation
pDC: plasmacytoid dendritic cell
LC: Langerhans Cell
cDC: classical dendritic cells
DTR: diphtheria toxin receptor
DT: diphtheria toxin
TCM: central memory T cells
TEM: effector memory T-cells
MPEC: memory-precursor effector cells
SLEC: short-lived effector cells

## Acknowledgements

We thank Paula Longhi for advice regarding dendritic cell depletion models and Richard Stanton for critical reading of the manuscript. This research was funded by a Wellcome Trust Senior Research Fellowship (Ref. 207503/Z/17/Z), a Wellcome Trust Collaborative Award in Science (Ref. 209213/Z/17/Z), an MRC Confidence in Concept Award, a Systems Immunity University Research Institute PhD Studentship and a Cardiff University School of Medicine PhD Studentship.

## Author Contributions

SD, SGB, MM, PS, LC and MC performed the experiments

SD, SGB and IH designed the experiments

SD and IH wrote the paper

## Supplemental Figure Legends

**Figure S1: Dendritic cell depletion in different DTR mouse strains**

**A:** transgenic BDCA2-DTR mice were injected i.p. with 300ng diphtheria toxin (+DT) or not (-DT) 24h before leukocytes were isolated from spleens and frequencies of plasmacytoid dendritic cells (pDCs) (B220^+^, SigLecH^+^) were quantified using flow cytometry. Data shown as mean ± SEM (n= 4 per group). *p ≤ 0.05. **B:** Transgenic LANG-DTREGFP mice were injected i.p. with 300ng diphtheria toxin (+DT) or not (-DT) 24h before DC migration assay was performed and Langerhans cells (LCs) (CD11c^+^ EPCAM1^+^) were quantified using flow cytometry. Data shown as mean ± SEM (n= 1-DT and 2 +DT). **C:** XCR1-DTR bone marrow chimeras were injected i.p. with 300ng diphtheria toxin (+DT) or not (-DT) on day −3, and day −1. Leukocytes were isolated from spleens on day +1 and frequencies as well as total numbers of professional cross-presenting DCs (CD11c^+^ ClassII^+^CD8^+^) were quantified using flow cytometry. Data shown as mean ± SEM (n= 1-DT and 4 +DT). **D:** zDC-DTR bone marrow chimeras were injected i.p. with 300ng diphtheria toxin (+DT) or not (-DT) 48h before leukocytes were isolated from spleens and frequencies as well as total numbers of classical dendritic cells (cDCs) (CD11c^+^ClassII^+^) were quantified using flow cytometry. Data shown as mean ± SEM (n= 1-DT and 2 +DT). Representative flow cytometry plots are shown for all DC subsets.

