## Supplementary figures and images for "Optimal CD8^+^ T-cell memory formation following subcutaneous cytomegalovirus infection requires virus replication but not early dendritic cell responses"

### Supplemental Figure 1

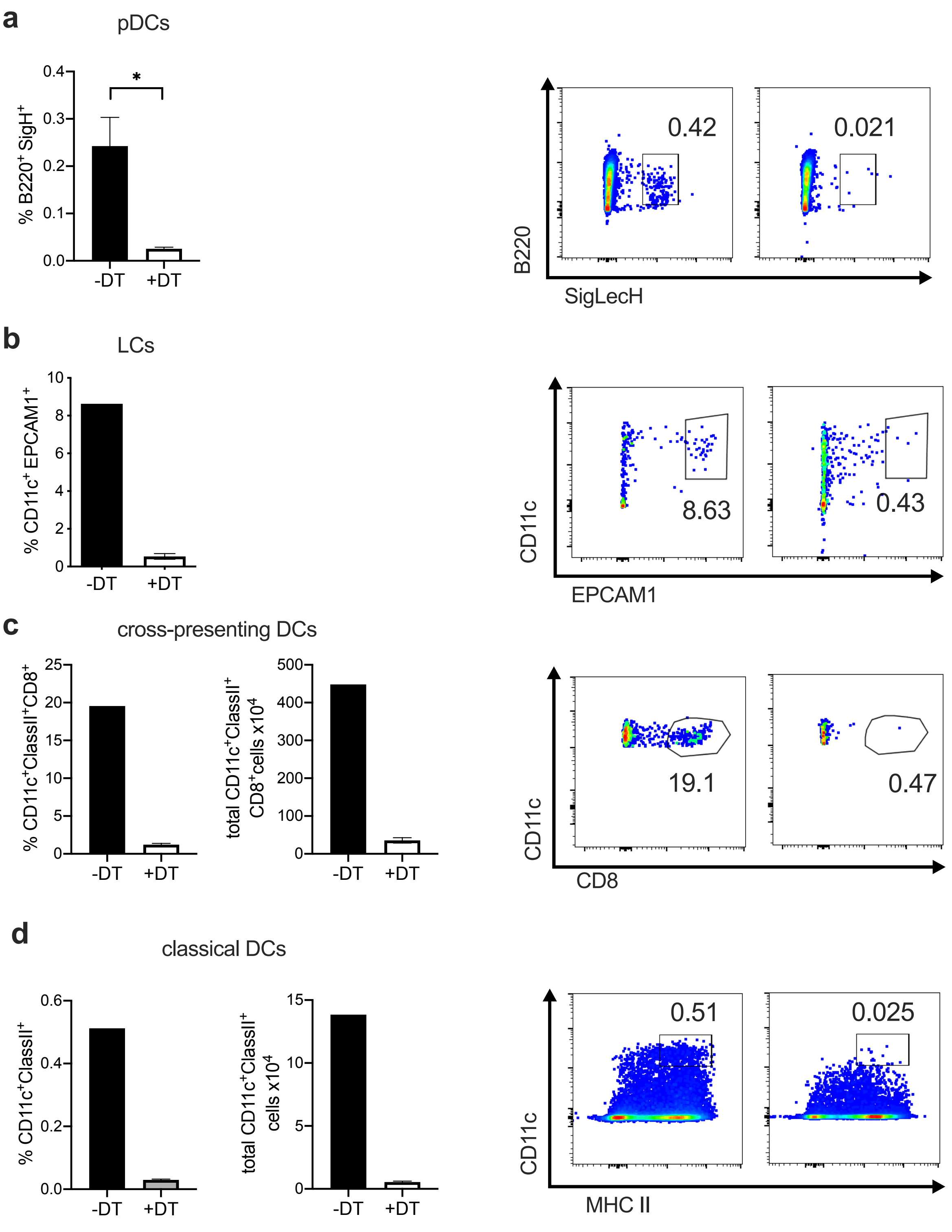
